# Systemic Treatment with Nicotinamide Riboside is Protective in Three Mouse Models of Retinal Degeneration

**DOI:** 10.1101/866798

**Authors:** Xian Zhang, Nathaniel F. Henneman, Preston E. Girardot, Jana T. Sellers, Micah A. Chrenek, Ying Li, Jiaxing Wang, Charles Brenner, John M. Nickerson, Jeffrey H. Boatright

## Abstract

Purpose: The retina is highly metabolically active, suggesting that metabolic dysfunction could underlie many retinal degenerative diseases. Nicotinamide adenine dinucleotide (NAD^+^) is a cofactor and a co-substrate in several cellular energetic metabolic pathways. Maintaining NAD^+^ levels may be therapeutic in retinal disease since retinal NAD^+^ levels decline with age and during retinal damage or degeneration. The purpose of this study was to investigate whether systemic treatment with nicotinamide riboside (NR), a NAD^+^ precursor, is protective in disparate models of retinal damage or degeneration.

**Methods:** Three mouse models of retinal degeneration were tested: an albino mouse model of light-induced retinal degeneration (LIRD) and two models of retinitis pigmentosa (RP), including a mouse line deficient in interphotoreceptor binding protein (IRBP) gene expression (IRBP KO), and a naturally-occuring cGMP phosphodiesterase 6b mutant mouse model of RP (the Pde6b^rd10^ mouse). Mice were intraperitoneally (IP) injected with PBS or NR at various times relative to damage or degeneration onset. One to two weeks later, retinal function was assessed by electroretinograms (ERGs) and retinal morphology was assessed by optical coherence tomography (OCT). Afterwards, retina sections were H&E stained for morphological analysis or by terminal deoxynucleiotidyl transferase dUTP nick and labeling (TUNEL). Retinal NAD^+^/NADH levels were enzymatically assayed.

**Results:** The retinal degeneration models exhibited significantly suppressed retinal function, and where examined, severely disrupted photoreceptor cell layer and significantly decreased numbers of nuclei and increased accumulation of DNA breaks as measured by TUNEL-labeled cells in the outer nuclear layer (ONL). These effects were prevented by various NR treatment regimens. IP treatment with NR also resulted in increased levels of NAD^+^ in retina.

**Conclusions:** This is the first study to report protective effects of NR treatment in mouse models of retinal degeneration. The positive outcomes in several models, coupled with human tolerance to NR dosing, suggest that maintaining retinal NAD^+^ via systemic NR treatment should be further explored for clinical relevance.

## Introduction

Retinal diseases are a leading cause of blindness globally, accounting for more than 25% of all vision impairment (over 55 million people).^1^ Macular degenerations such as age-related macular degeneration (AMD), inherited retinal degenerations such as retinitis pigmentosa (RP), and several other retinopathies, syndromes, and dystrophies are linked to mutations in over 300 different genes or loci (Retnet; sph.uth.edu/retnet/disease.htm; update 10/29/2019). In addition to heterogeneous heritable risk factors, several retinal diseases such as AMD have strong environmental risk factors such as smoking, or the case of retinal detachments with degenerative outcome, extreme myopia.^2–5^ Although treatment strategies based on specific genetic mutations, risk factors, or combinations thereof are being developed (one gene therapy study has already entered the clinic^6^), approaches with targets that are common to many diseases may have greater impact.

Though of varied etiology, the outcome that is common to retinal degenerative diseases is the progressive loss of rod and cone photoreceptor cells and retinal pigment epithelial (RPE) cells. The retina is the most metabolically demanding tissue in the body, and experiences significant, persistent oxidative stress arising from its primary function, visual transduction.^7, 8^ Photoreceptor cells in particular are highly metabolically active as their normal function includes maintenance of the “dark current” and continuous production of proteins that support their specialized function,^8^ many of which are lost daily due to outer segment disc shedding.^9^ Because of this, photoreceptors are thought to be especially sensitive to dysregulation of bioenergetic metabolism.^10–12^ Indeed, genetic mutations that degrade photoreceptor bioenergetic metabolism cause blindness, including mutations of Krebs cycle enzymes that cause forms of RP^13^ and mutations in mitochondrial DNA (mtDNA) that cause Leber Hereditary Optic Neuropathy (LHON).^14^ Also, as photoreceptor and RPE cells exist within a metabolic symbiosis,^15^ functional compromise in one cell type may negatively affect function in the other.^10^ Thus, targeting metabolic pathways of photoreceptor or RPE cells may be therapeutic across a variety of retinal diseases.

Nicotinamide adenine dinucleotide (NAD^+^) is a cofactor in glycolysis and the Krebs cycle and a co-substrate for NAD^+^-consuming enzymes.^8, 16^ NAD^+^ levels decline with age and in neurodegeneration, possibly in response to increased activity of NAD^+^-consuming enzymes involved in DNA repair, such as poly (ADP-ribose) polymerase (PARP).^16^ Maintenance of NAD^+^ levels is critical to retinal health. Mutations in nicotinamide mononucleotide adenylyltransferase-1 (NMNAT1), a NAD^+^ salvage pathway enzyme, causes Leber Congenital Amaurosis Type 9 (LCA9).^17–19^ Experimentally inhibiting nicotinamide phosphoribosyltransferase (NAMPT), a key enzyme in a NAD^+^ salvage biosynthesis pathway, with pharmacological agents causes retinal degeneration.^20^ In a seminal series of experiments, Apte and colleagues established the importance of NAD^+^ in retinal health and the role of NAD^+^ deficiency in retinal degeneration.^8^ They found that retinal NAD^+^ levels decline in models of retinal degeneration and that mice made deficient in NAMPT have diminished retinal NAD^+^ and develop retinal degeneration.^21^ Further, they demonstrated that systemic treatment with the NAD^+^ precursor nicotinamide mononucleotide (NMN) was protective in mouse models of retinal degeneration.^21^ Similarly, John and colleagues have demonstrated that NAD^+^ is critical to retinal ganglion cell (RGC) health, that NAD^+^ levels decline in models of glaucoma, and that treatment of these models with NAM maintains or increases NAD^+^ in RGCs via the NAMPT pathway, and that this protects function and optic nerve morphology.^22, 23^

Brenner and colleagues discovered an alternative NAD^+^ salvage biosynthesis pathway in which nicotinamide riboside (NR), a form of vitamin B3 that is found in milk and other foods,^24, 25^ can be taken up from oral dosing in humans.^26^ NR enters cells through nucleotide transporters and is then converted to NMN then to NAD^+^ by NR kinases (NMRK1 and NMRK2) and NMNAT isozymes.^27, 28^ They and others find that administration of NR to rodents prevents cognitive decline and amyloid-beta peptide aggregation,^29^ prevents noise-induced neurite degeneration and hearing loss,^30^ protects in several models of neuropathies,^31, 32^ and protects against excitotoxicity-induced or axotomy-induced axonal degeneration.^33, 34^ The NMRK kinases are expressed in many neuronal subtypes, and may be upregulated following neuronal injury or during extreme energetic stress.^16^ NR treatment also protects in neurodegeneration models. NR treatment increases NAD^+^ and protects mitochondrial function in cultured neurons from Parkinson Disease (PD) patients and prevents age-related dopaminergic neuronal loss and motor decline in Drosophila models of PD.^35^ In a new mouse model of Alzheimer Disease (AD), NR provided in drinking water improved learning, memory, and motor function. In addition, NR supplementation led to normalized NAD^+^/NADH ratios, better synaptic transmission, and decreased DNA damage.^36^

Given this background supporting a neuroprotective role of NR in several neuronal degenerations, we hypothesized that NR would be effective in more than one retinal degeneration model including both inherited retinal degenerations and experimentally induced retinal degenerations. In this study, we tested whether systemic delivery of NR is protective in three disparate mouse models of retinal degeneration: a mouse model of light-induced retinal degeneration (LIRD),^37–39^ and two models of autosomal recessive retinitis pigmentosa (arRP), the IRBP knock-out (KO) mouse^40, 41^ and the rd10 mouse.^42^ NR treatment increased retinal NAD^+^ and the NAD^+^/NADH ratio and protected retinal morphology and function in all three models. This is the first study to report protective effects of NR treatment in models of retinal degeneration and suggests that maintaining retinal NAD^+^ via systemic NR treatment should be further explored for clinical relevance.

## Materials and Methods

### Animal models

All mouse procedures were approved by the Emory Institutional Animal Care and Use Committee and followed the ARVO Statement for the Use of Animals in Ophthalmic and Vision Research. For the LIRD model, adult (3 months old) male Balb/c mice were obtained from Charles River Laboratory (Wilmington, MA, USA) and were housed under a 12:12-hour light-dark cycle (7 AM on and 7 PM off). During the light cycle, light levels measured at the bottom of mouse cages ranged from 5 to 45 lux. Induction of LIRD was previously described.^37–39^ Briefly, Balb/c mice were placed individually into white, opaque, standard-sized and -shaped housing cages. LED light panels were placed on top of the cages. Mice were exposed to 3,000 lux light for 4 hours. After this exposure, mice were returned to home cages under normal lighting conditions for the remainder of the experiment. Mice had access to standard mouse chow (Lab Diet 5001; LabDiet, Inc., St. Louis, MO, USA) ad libitum throughout the study except during bright light exposure. At the time of light induction they weighed 22 to 28 g. NR treatment did not significantly alter weight (data not shown). Mice were euthanized by asphyxiation with CO_2_ gas for all experiments.

IRBP KO mice^43^ used in these experiments were originally created from 129/Ola embryonic stem cells and were backcrossed against C57BL/6J for 10 generations.^44^ Food and water were provided ad libitum, with 12:12 light-dark cycling. Genotypes were verified with specific primers^45^ in all strains. The ERG phenotype in the IRBP KO mouse was reduced by 40% of the wild type (WT) a-wave magnitude at postnatal 30 (P30) as previously shown,^43^ validating use of the current KO mice in this study. Male and female mice from each litter were randomly divided at P18 to receive NR treatment (1000 mg/kg) or PBS.

rd10 mice on a C57BL/6J background^46, 47^ were obtained from Jackson Laboratories (Bar Harbor, Maine). In the rd10 retina, rod cell death begins at about P15, and by P25 almost all of the rod photoreceptors have been lost.^48^ In our experiments, male and female mice from each litter were randomly divided at P6 to receive subcutaneous NR (1000 mg/kg) or PBS treatment until P14 and then received IP NR or PBS treatment until P27.

### Drug Administration

NR was kindly provided by Dr. Charles Brenner (ChromaDex, Item #ASB-00014332-101, Lot# 40C910-18209-21). NR and vehicle (phosphate buffered saline, PBS, VWRVK813, Cat#97063-660, 1X solution composition:137 mM NaCl, 2.7 mM KCl, 9.5 mM phosphate buffer.) Solutions were made fresh each day. Balb/c mice received IP injections of either PBS alone or NR (1000 mg/kg dose in PBS; specified by experiment as reported in Results) using an injection volume of 10 μl of solution per gram of mouse body weight in accordance with in vivo rodent experiments of Yen et al.^49^ For LIRD experiments, two injections were administered before a toxic light exposure. One injection was performed the day before (at 4 pm), and the other injection was performed at 9 am the morning of light insult. Toxic light exposure was from 10 am to 2 pm. The IRBP KO mice received treatment 5 days per week from P18 to P33. rd10 mice were injected 5 days per week from P6 to P27. Injections from P6-P14 were subcutaneous, and from P15-P27 were IP.

### Electroretinograms (ERG)

The complete ERG protocol was previously detailed.^50^ Briefly, mice were dark-adapted overnight. In preparation for ERGs, mice were anesthetized with IP injections of ketamine (10 mg/ml: AmTech Group Inc.) and xylazine (100 mg/ml; AKORN Animal Health, Lake Forest, IL).^50^ Proparacaine (1%; AKORN, Lake Forest, IL) and tropicamide (1%; AKORN, Lake Forest, IL) eye drops were administered to reduce eye sensitivity and dilate pupils. Once anesthetized, mice were placed on a heating pad inside a Faraday cage in front of a UBA-4200 Series desktop Ganzfeld stimulator (LKC Technologies, MDIT-100). A DTL fiber active electrode was placed on top of each cornea. A drop of Refresh Tears (Allergan, Dublin, Ireland) was added to each eye to maintain conductivity with the electrode fibers. The reference electrodes was a 1-cm needle inserted into the cheeks, and the ground electrode placed in the tail. ERGs were recorded for the scotopic condition (0.00039-25.3 cd s/m^2^ with increasing flash stimulus intervals from 2 to 62.6 s). Mice recovered from anesthesia individually in cages placed partly on top of heated water pads (39 **°**C). For the LIRD model, ERGs were performed 1 week after toxic light exposure and again at 2 weeks following light exposure. As for the two inherited RP models, ERGs were performed after 10 or 20 injections of NR.

### In vivo ocular imaging

Spectral domain optical coherence tomography (SD-OCT) was conducted immediately after ERG measurement, when mice were still anesthetized and their pupils were still dilated. A Micron IV SD-OCT system with fundus camera (Phoenix Research Labs, Pleasanton, CA) and a Heidelberg Spectralis HRA+OCT instrument with +25D lens (Heidelberg Engineering, Heidelberg, Germany) were used in tandem sequentially to assess ocular posterior segment morphology in section and *en face*. Using the Micron IV system, image-guided OCT images were obtained for the left and right eyes after a sharp and clear image of the fundus (with the optic nerve centered) was obtained. SD-OCT imaging was a circular scan about 100 μm from the optic nerve head. Fifty scans were averaged. The retinal layers were identified according to published nomenclature.^51^ Total retinal thickness and thickness of the individual retinal layers were analyzed using Photoshop CS6 (Adobe Systems Inc., San Jose, CA). The number of pixels was converted into micrometers by multiplying by the micrometers/pixel conversion factor (1.3 microns = 1 pixel). Immediately after imaging on the Micron IV system, a rigid contact lens was placed on the eye (Back Optic Zone Radius: 1.7 mm, diameter: 3.2 mm, power: PLANO) and blue autofluorescence imaging at the layer of the photoreceptor-RPE interface was conducted using the Heidelberg Spectralis HRA+OCT instrument. During imaging and afterwards through anesthesia recovery, mice were kept on a water circulating heat pad set to 39 **°**C to maintain body temperature.

#### Histology and Morphometrics

The Balb/c mice were sacrificed the same day after in-vivo measurments. We followed the recommendations of Howland and Howland^52^ for nomenclature of axes and planes in the mouse eye, as indicated in Figure 1. Histologic and morphometric procedures followed standard techniques.^46, 53^ Eyes were dehydrated, embedded in paraffin, and sectioned through the sagittal plane on a microtome at 5 µm increments. Sections were cut on a vertical (sagittal) plane through the optic nerve head (ONH) and the center of the cornea to ensure that consistent regions were examined between animals. Slides were de-paraffinized across five Coplin jars with 100 ml of xylene for two minutes each, consecutively. Then slides were rehydrated in a series of 100 ml ethanol solutions for two minutes each: 100%, 90%, 80%, 70%, 60%, 50%. Slides were immersed in PBS for five minutes twice. After rehydration, a TUNEL assay was performed on some sections according to the protocol for the DeadEnd Fluorometric TUNEL Kit (Promega, Fitchburg, WI). Stained sections were imaged using fluorescent microscopy and TUNEL-positive cells in the outer-nuclear layer (ONL) were manually counted for each whole retina using Adobe Photoshop Creative Suite 6 (Adobe Systems Inc. San Jose, CA). Some sections were used for hematoxylin and eosin (H&E) staining. ONL nuclei were counted within 100 μm-wide segments spaced at 250, 750, 1250, 1750 μm from the optic nerve head in both the inferior and superior directions. Mean counts of n = 3-6 retinas per group were plotted as a spidergram diagram.

**Figure 1.**
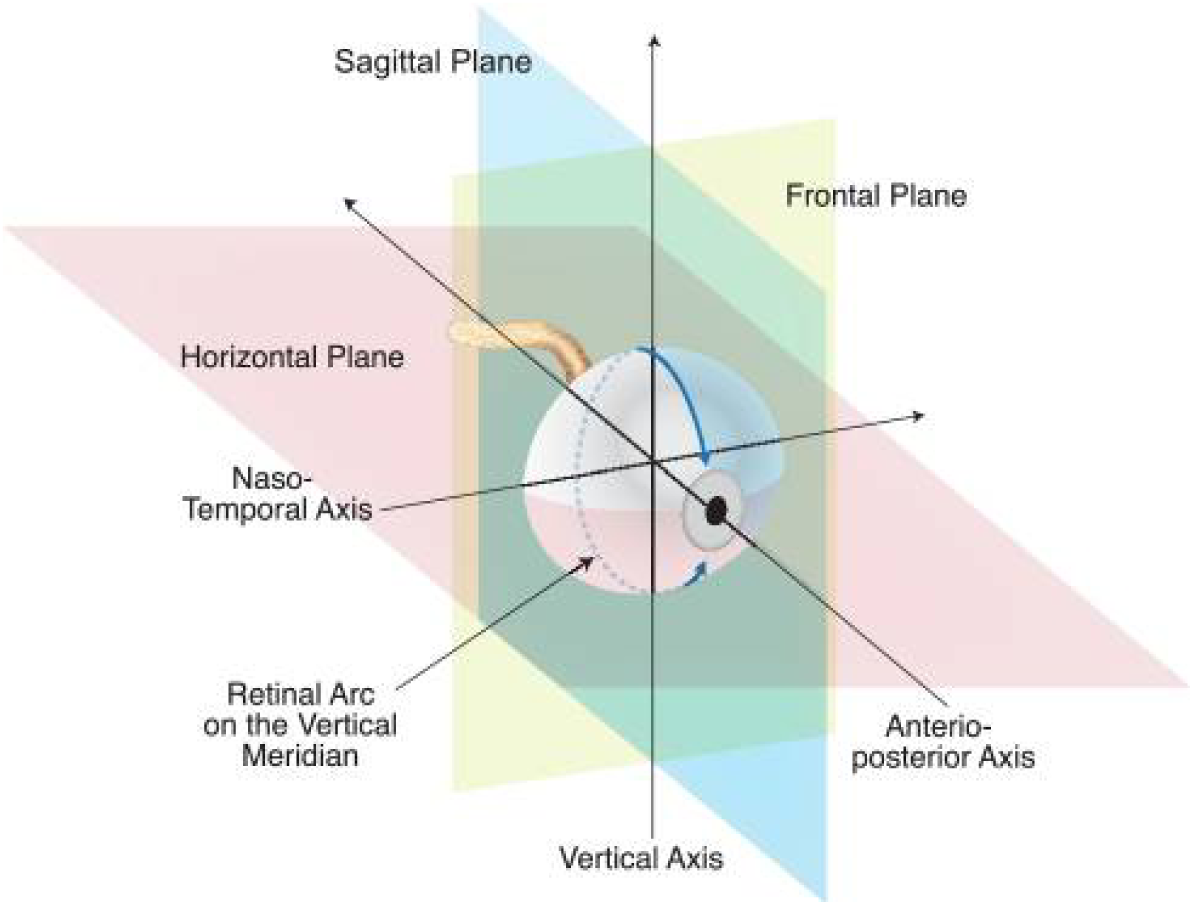
Naming conventions recommended by Howland and Howland for planes and axes of the vertebrate eye regardless of species. A histologic section cut on the vertical plane through the great meridian is illustrated.^40^ The horizontal, vertical, and frontal planes are marked, as are the anterior-posterior (A-P) axis (also known as optic axis), the nasal-temporal (N-T) axis, and the superior-inferior (S-I; vertical) axis. Republished from Wisard at el. 2011 (citation 40), with permission.

#### Histology and Morphometrics

NAD^+^/NADH measurements

Levels of NAD^+^ in retina homogenates were measured using a commercially available kit by following manufacturer’s instructions (Abcam; ab. 65348; #Lot: GR3226737-3; San Francisco, CA). In brief, retina samples were homogenized in extraction buffer. Extracted samples were separated in two aliquots. One was used to measure total NAD (NADt). The other was heated to 60℃ for 30 minutes to convert NAD^+^ to NADH. Samples were placed in a 96-well plate. Then, NADH developer was added into each well and incubated at room temperature for 1-4 hours. The plate was placed into a hybrid reader and read every half hour at OD 450nm while the color was still developing. We used the 2-hour data, at which the reaction was well developed. NADt and NADH concentration was quantified comparing with NADH standard curve data. In the end, NAD^+^ was calculated with the equation NAD^+^=NADT-NADH.

### Statistical Analyses

Statistical analyses were conducted using Prism 8.1.1 Software (GraphPad Software Inc. La Jolla, CA, USA). One-way ANOVA with Tukey’s post-hoc test, two-way ANOVA with Sidak’s post-hoc test, and Student’s *t-*tests were performed for ERG, biochemical, and morphometric data. Tukey post-hoc test is considered to be more powerful in comparing every mean with every other mean, and so is most appropriate for our use of the one-way ANOVA. Sidak’s test is more generalized, allowing for comparisons of just subsets of means, and thus recommended for our use of the two-way ANOVA. For all analyses, results were considered statistically significant if *p < 0.05*. All graphs display data as mean ± SEM. The stated *n* is the number of animals or individual eye used in each group.

## Results

### NR treatment preserves retina function in three mouse models of retinal degeneration

ERG a- and b-wave mean amplitudes of PBS-treated Balb/c mice were significantly reduced one week after exposure of to 3,000 lux light for 4 hours. Representative ERG waveforms of one eye of a single mouse from each treatment group are shown in Fig. 2. This functional loss was entirely prevented in mice treated with NR (Fig. 3A, 3B). Functional loss was diminished significantly, but not entirely, in the two genetic mouse models of RP. The IRBP KO mouse degeneration starts around P24 with a burst of apoptosis in the outer nuclear layer, followed by a very gradual loss of photoreceptor cells over the lifespan of the animals.^40^ This gradual loss is accompanied by a gradual decline of ERG waveform amplitudes.^54^ Scotopic and photopic ERG a- and b-wave amplitudes were greater following daily NR versus PBS injections (Fig. 4). The rd10 mouse has an early onset and rapid photoreceptor degeneration due to a mutation in the beta subunit of rod phosphodiesterase, with significant morphological and functional losses apparent at P16 that are nearly complete by P30.^42, 50^ Daily IP injections of NR significantly preserved scotopic ERG b-wave amplitudes and photopic ERG a- and b-wave amplitudes (Fig 5). This protective effect persisted at P38, 10 days after treatment had been halted (Fig 6). These data suggest that NR treatment protects against of retinal damage and degeneration.

**Figure 2.**
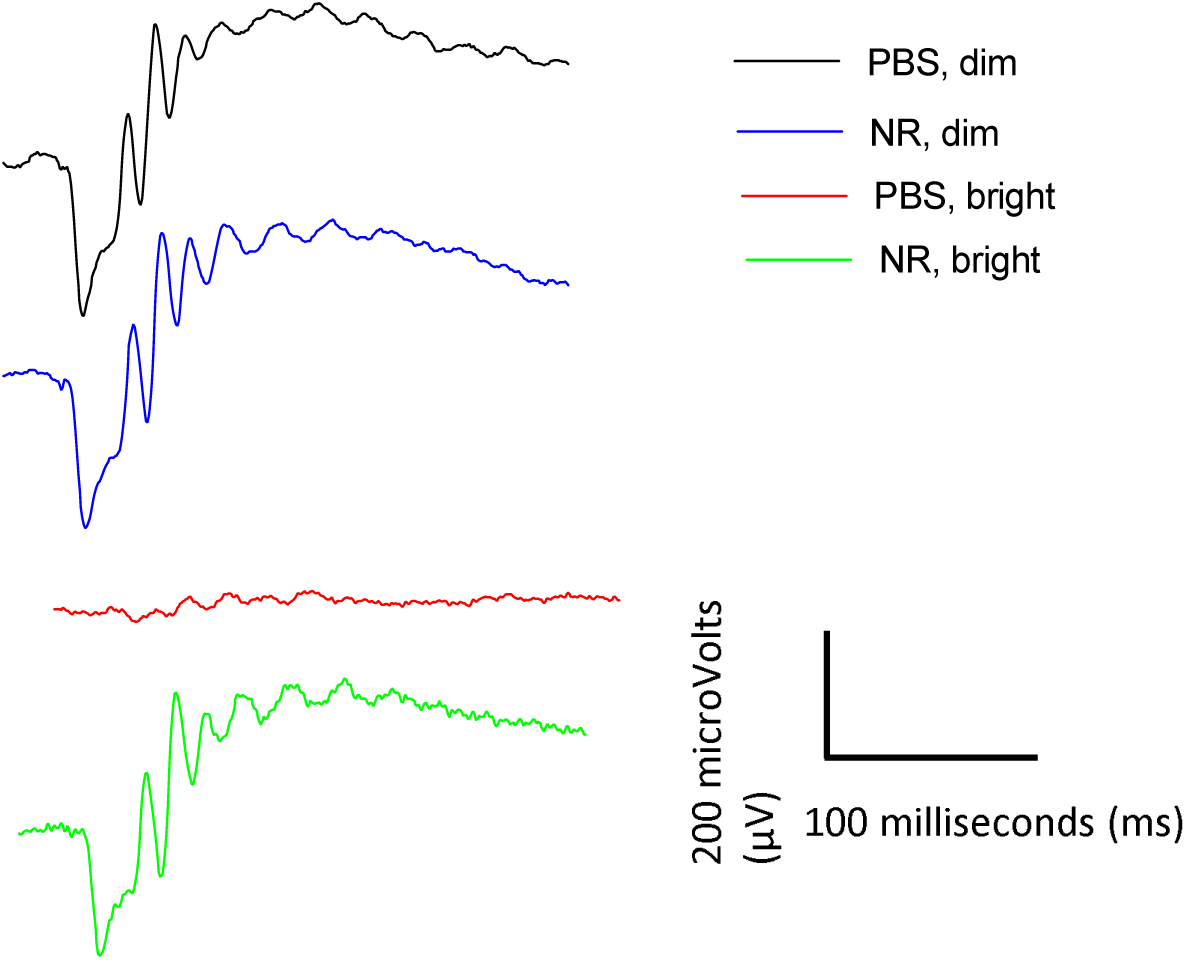
NR (1000 mg/kg) treatment preserves a- and b-wave amplitudes 1 week following toxic light exposure. Representative ERG waveforms of one eye of a single mouse from each treatment group. Mice treated with PBS and exposed to 3,000 lux for 4 hours show the most diminished a- and b-wave amplitudes. Treatment with NR protected against the loss of a- and b-wave amplitudes. The ERG flash intensity was 24.9 cd-s/m^2^ and was the same for all four mice.

**Figure 3.**
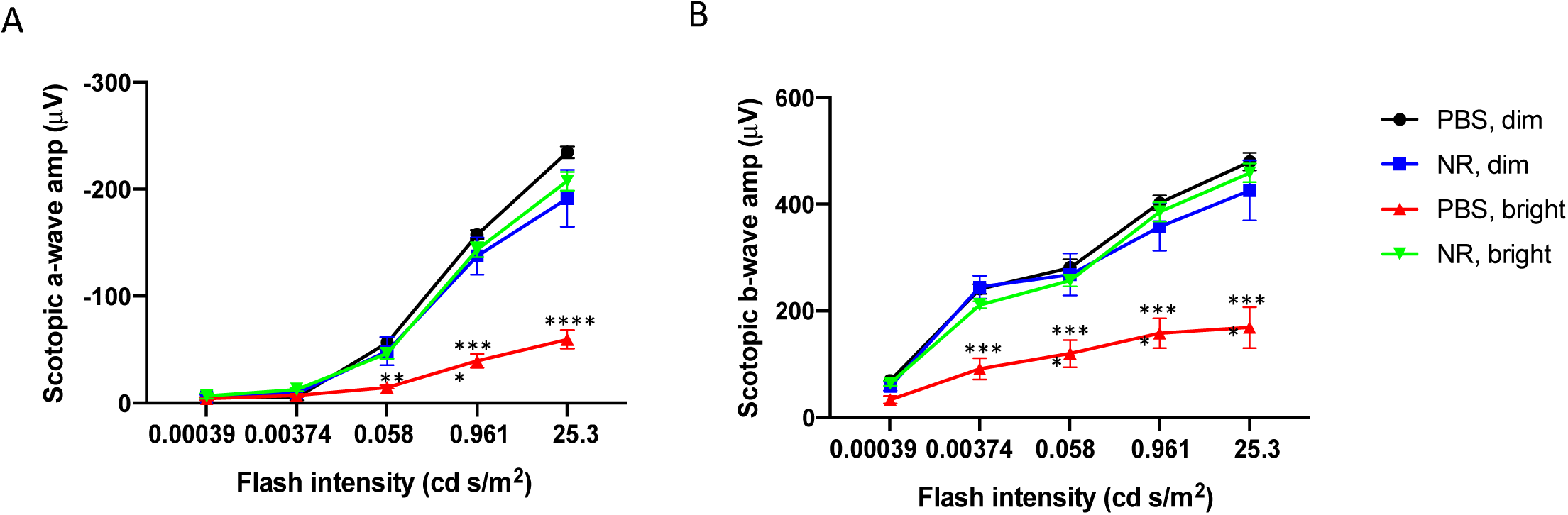
NR treatment preserves retinal function in LIRD mouse. Scotopic ERG a-wave (A) and b-wave (B) mean amplitudes from LIRD mice at 1 week after degeneration induction. Mice treated with PBS and exposed to 3,000-lux light for 4 hours to induce degeneration exhibited a- and b-wave mean amplitudes (red) that were statistically significantly diminished compared to those of dim groups (blue and black). However, the mean ERG amplitudes of mice undergoing retinal degeneration and treated with NR (green) were statistically indistinguishable from those treated with PBS and exposed to bright light. **P < 0.01, ***P < 0.001, ****P < 0.0001 versus other groups by two-way ANOVA with Tukey’s multiple comparisons test. n=8-12 eyes per group. Error bars represent SEM.

**Figure 4.**
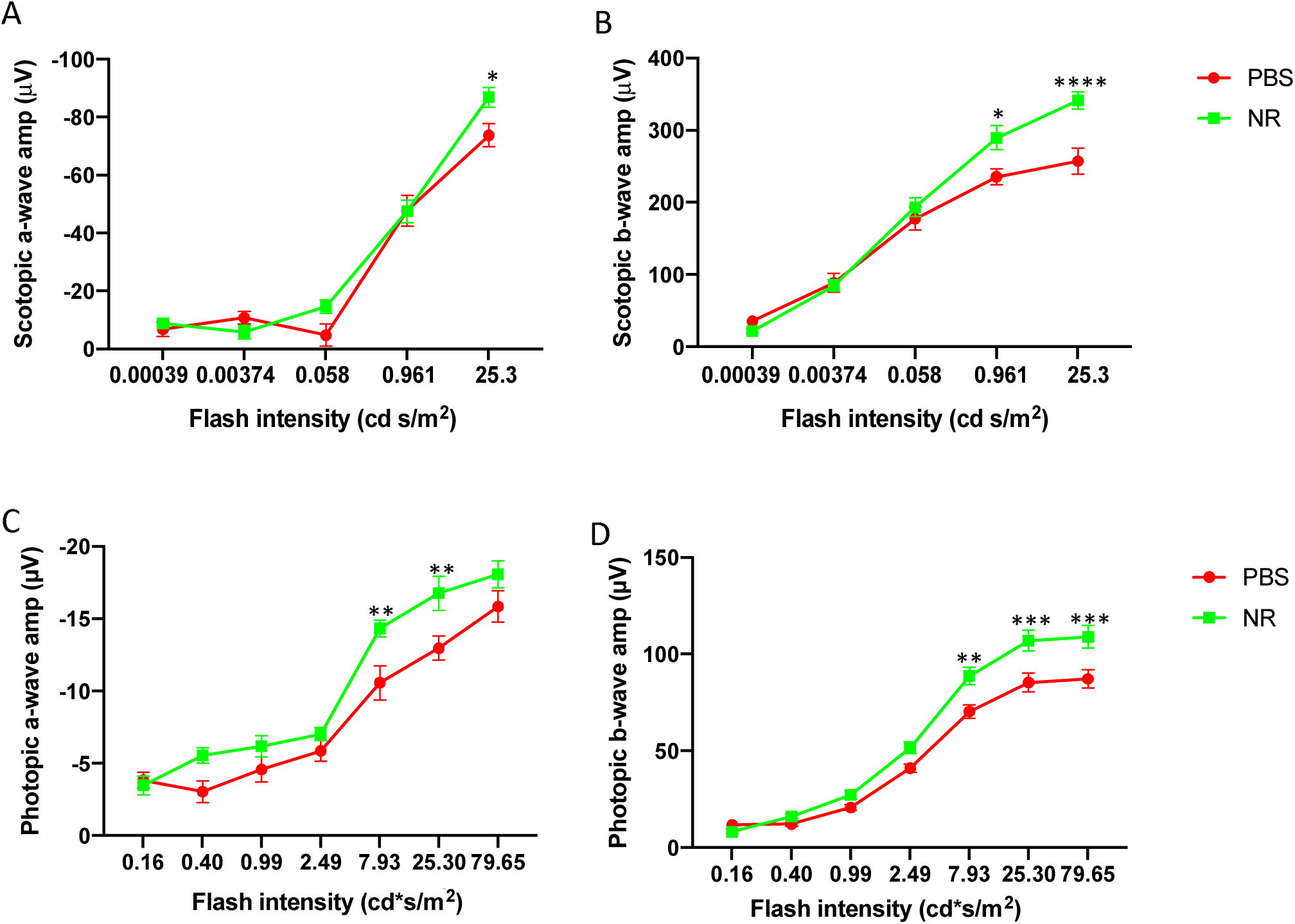
NR treatment preserves retinal function in IRBP KO mice. Scotopic ERG a-wave (A) and b-wave (B) and photopic a-wave (C) and b-wave (D) mean amplitudes from IRBP KO mice which were IP injected 5 days a week from P18 to P33. The mean ERG amplitudes of mice treated with NR (green) were statistically greater than those treated with PBS. *P<0.05, **P < 0.01, ***P < 0.001, ****P < 0.0001 by two-way ANOVA with Sidak’s multiple comparisons non-parametric test; For each group, n=10-12 eyes. Error bars represent SEM.

**Figure 5.**
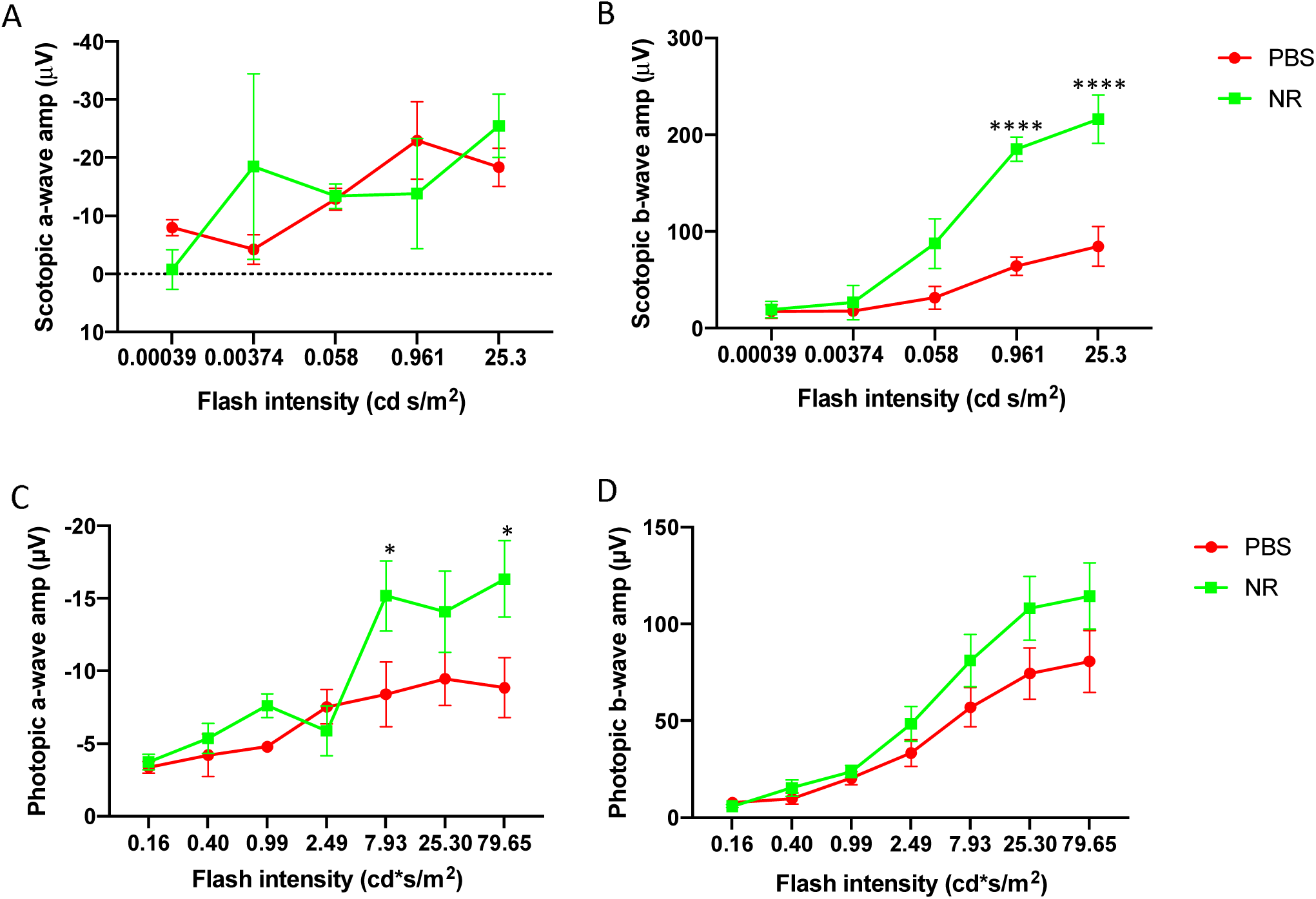
NR treatement preserves retinal function in PDE6brd10 mice at P28. Scotopic ERG a-wave (A) and b-wave (B) and photopic a-wave (C) and b-wave (D) mean amplitudes from rd10 mice which were IP injected 5 days a week from P6 to P28. The mean ERG amplitudes of rd10 mice treated with NR (green) were statistically greater than those treated with PBS. *P<0.05, ****P < 0.0001 by two-way ANOVA with Sidak’s multiple comparisons non-parametric test; For each group, n=6 eyes. Error bars represent SEM.

**Figure 6.**
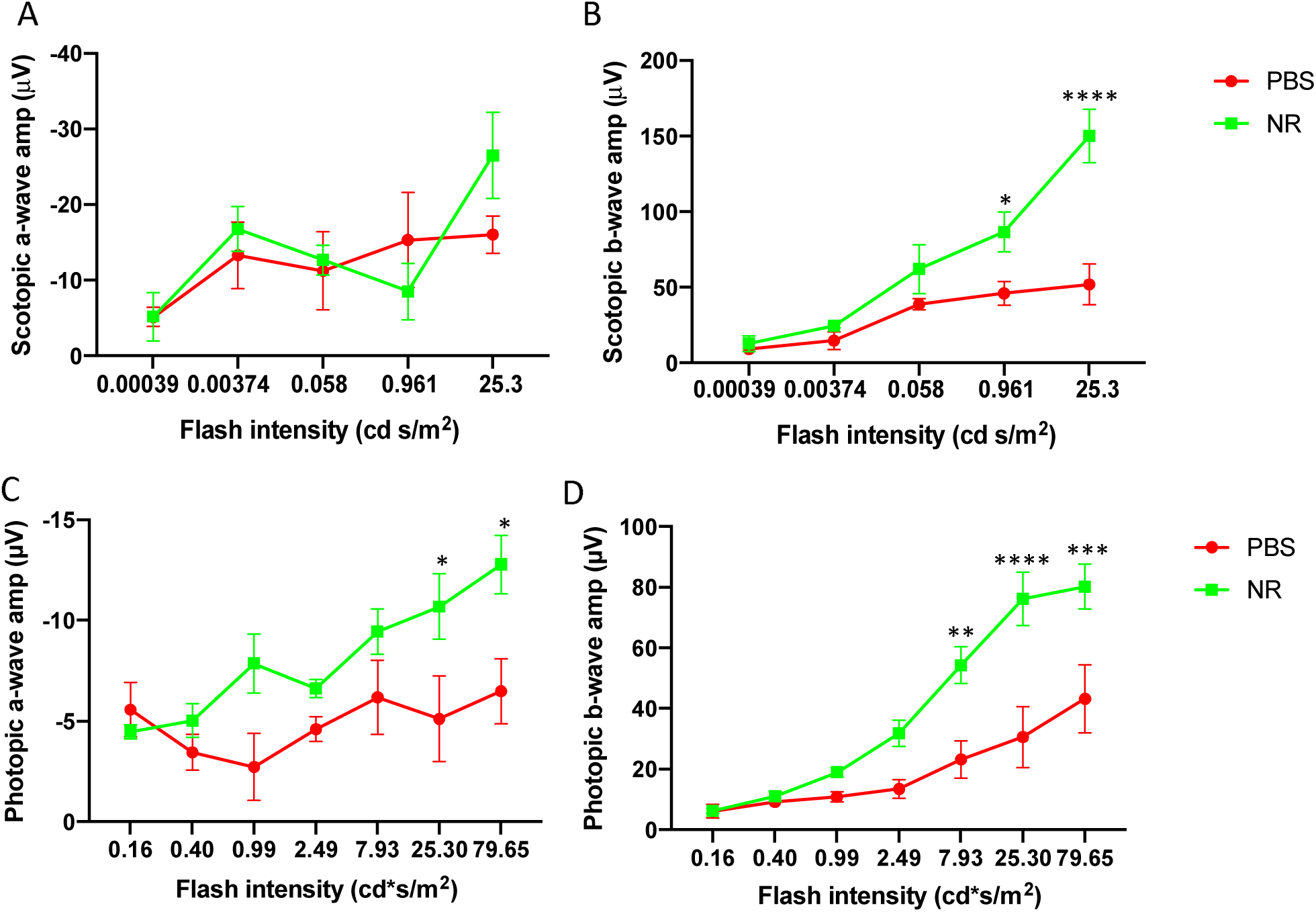
NR preserves retinal function in PDE6b^rd10^ mice P38 (10 days after cessation of treatment). Scotopic ERG a-wave (A) and b-wave (B) and photopic a-wave (C) and b-wave (D) mean amplitudes from rd10 mice at P38, following 21 treatments ending 10 day prior (P28). The mean ERG amplitudes of rd10 mice treated with NR (green) were statistically greater than those treated with PBS. *P<0.05, **P < 0.01, ***P < 0.001, ****P < 0.0001 by two-way ANOVA with Sidak’s multiple comparisons non-parametric test; For each group, n=6 eyes. Error bars represent SEM.

### NR treatment preserves retinal morphology in LIRD mouse

As imaged *in vivo* by fundus photography and SD-OCT one week after degeneration induction, the retinas of LIRD mice treated with PBS exhibited significant damage (Fig. 7). The representative fundus image of Fig. 7A shows the region being measured. Corresponding SD-OCT images show considerable thinning of the photoreceptor layer that is prevented by NR treatment (Fig. 7B). Quantification of total retina and photoreceptor layer thicknesses show that mice undergoing LIRD had significantly thinner retinas compared to non-induced mice, largely due to nearly complete loss of the photoreceptor layer. This was entirely prevented with NR treatment (Fig. 7C, 7D).

**Figure 7.**
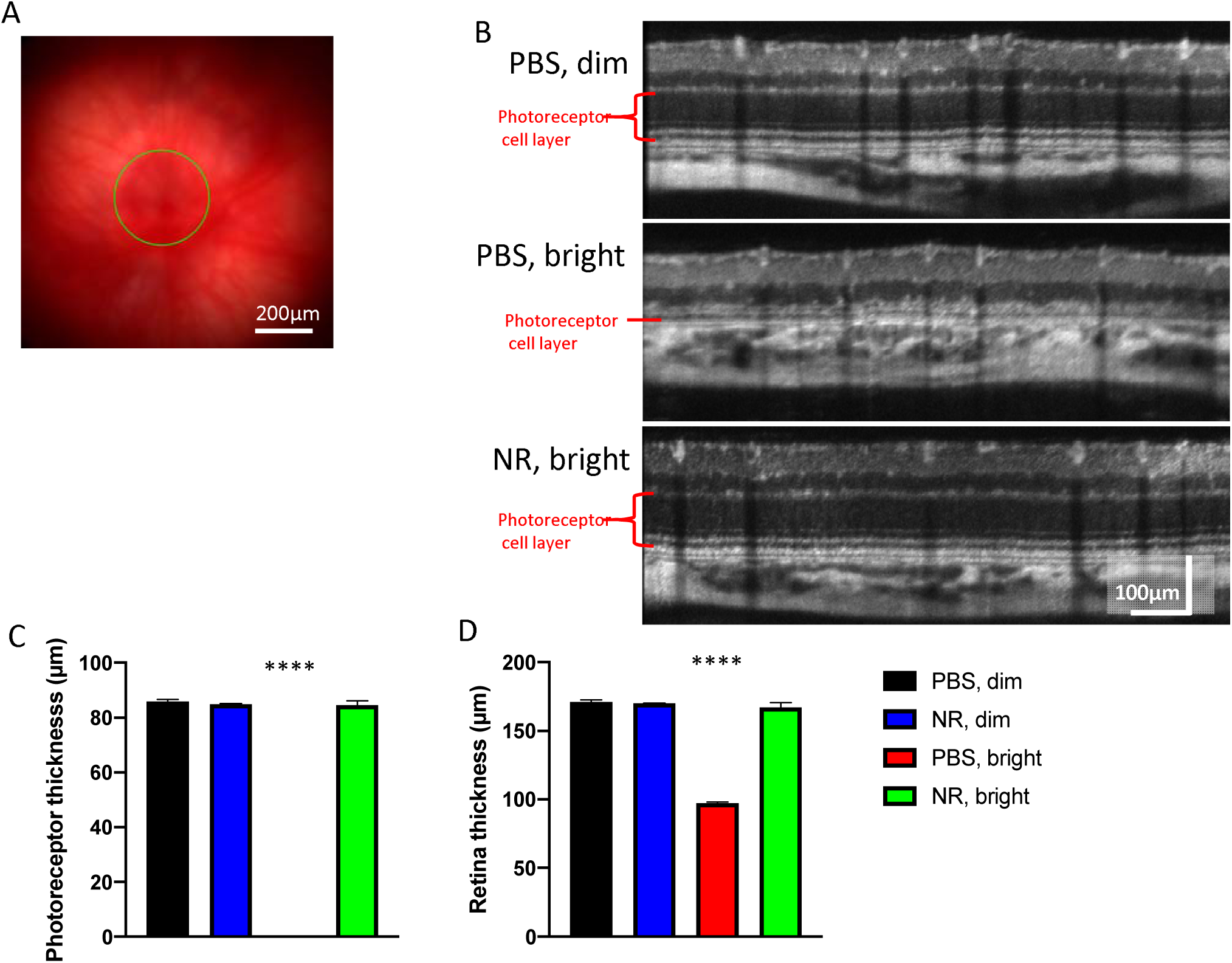
NR treatment preserves photoreceptor layer thickness and total retinal thickness as assessed in vivo following light-induced retinal degeneration. (A). Representative fundus image (B). OCT images from each group. The OCT image is a circular scan about 100 μm from the optic nerve head. Photoreceptor thickness (C) and retinal thickness (D) from Balb/c mice at one week after degeneration induction. Mice treated with NR and exposed to 3,000-lux light for 4 hours (red bar) exhibited great losses in thickness of the photoreceptor and retina layers, whereas induced mice treated with NR (green bar) exhibited statistically significant preservation of layer thickness. ****p<0.0001 versus all other group means; one-way ANOVA with Tukey’s multiple comparisons non-parametric test. n = 3-6 mice/group. Error bars represent SEM. Size marker represents 200 µm.

Photomicroscopy of H&E-stained sections of eyes harvested one week after LIRD induction showed marked degradation of morphology in the outer retina in LIRD mice treated with PBS compared to non-induced mice (Fig.8A, 8B, and Fig. S1). In induced mice treated with PBS, photoreceptor cell inner and outer segments and most of the nuclei of the ONL were eliminated, with loss predominantly centrally, (Fig. 8A-D, and Fig. S1). Nearly all of this degeneration was prevented in mice treated with NR (Fig. 8C and Fig. S1). Quantification of ONL nuclei counts across retinal sections confirmed significant losses due to degeneration in PBS-treated mice and confirmed nearly-complete preservation, in NR-treated mice (Fig. 8D and Fig. S1).

**Figure 8.**
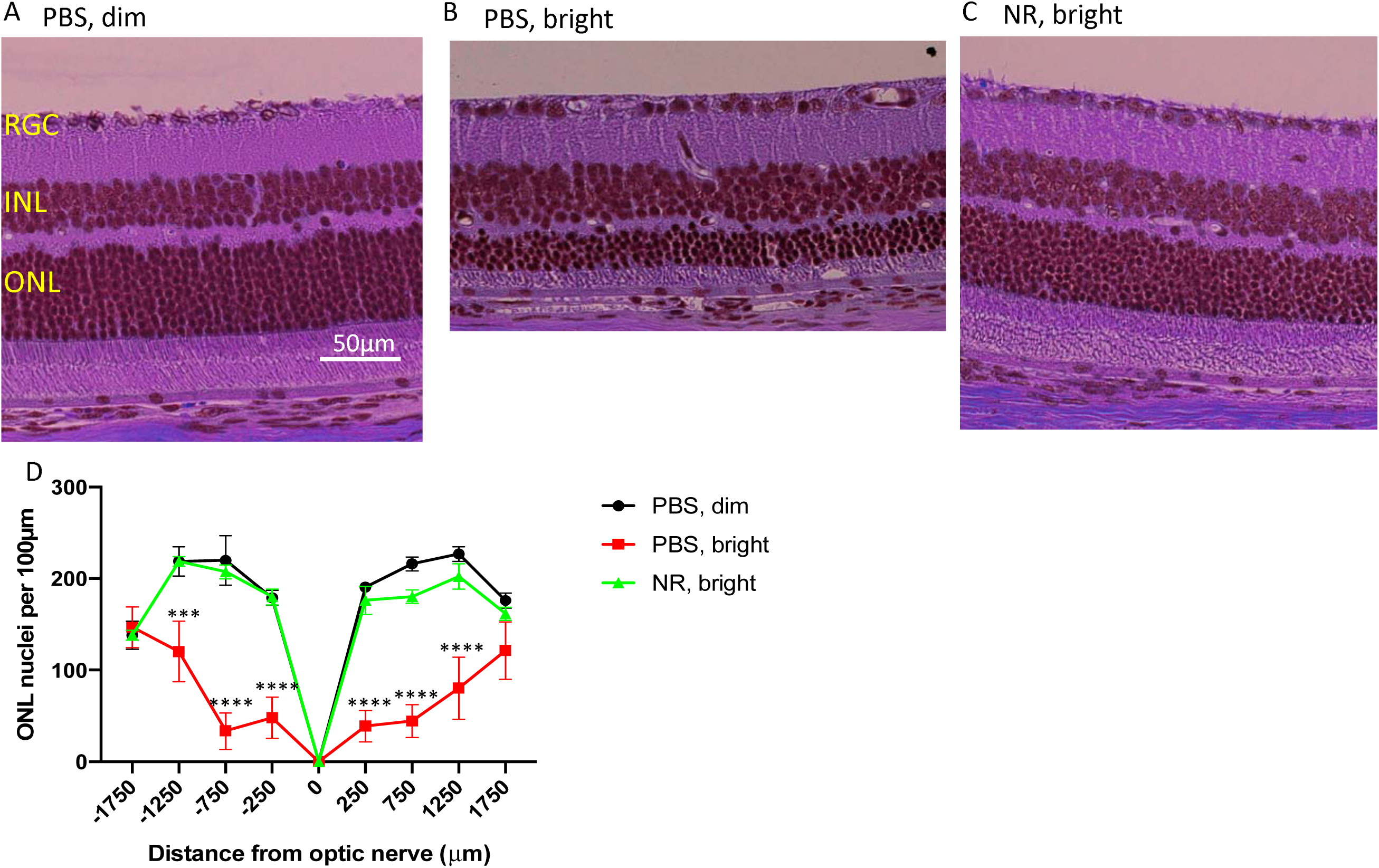
NR treatment preserves nuclei of the outer nuclear layer in the LIRD mice. (A)-(C). Representative H&E images of retina sections from each group at the region of 250-750 µm from optic nerve. Complete sections are shown in Supplemental Figure S1. (D). One week after degeneration induction, nuclei were counted in eight discrete regions of retinal sections starting at 250 µm from the optic nerve head and extending every 500 µm outward along both the dorsal/superior (positive values on abscissa) and ventral periphery/inferior (negative numbers on abscissa). ‘PBS, bright’ treated mice (red) showed significant loss of nuclei at six distances from the optic nerve head compared to the control group (black). However, NR treated mice (green), exhibited mean nuclei counts statistically indistinguishable from that of the control group (black) throughout the length of the retina. ***P < 0.001, ****P < 0.0001 versus other groups by two-way ANOVA with Tukey’s multiple comparisons non-parametric test. n = 3-6 retinal images/group. Error bars represent SEM. Size marker represents 50 µm.

### NR treatment prevents accumulation of TUNEL signal in photoreceptor cells of LIRD mice

Paraffin-embedded ocular sections from mice euthanized a week after toxic light induction were stained for TUNEL and DAPI to label nuclei that contained double-stranded DNA breaks (a marker of programmed cell death [apoptosis] or other forms of cell death).^55^ ONL TUNEL signal was high in retinas from LIRD mice treated with PBS compared to retinas from uninduced mice (Fig. 9A, 9B and Fig. S2). Induced mice treated with NR exhibited significantly less TUNEL signal (Fig. 9C and Fig. S2). These data suggest that NR treatment diminished or delayed apoptosis in photoreceptor cells.

**Figure 9.**
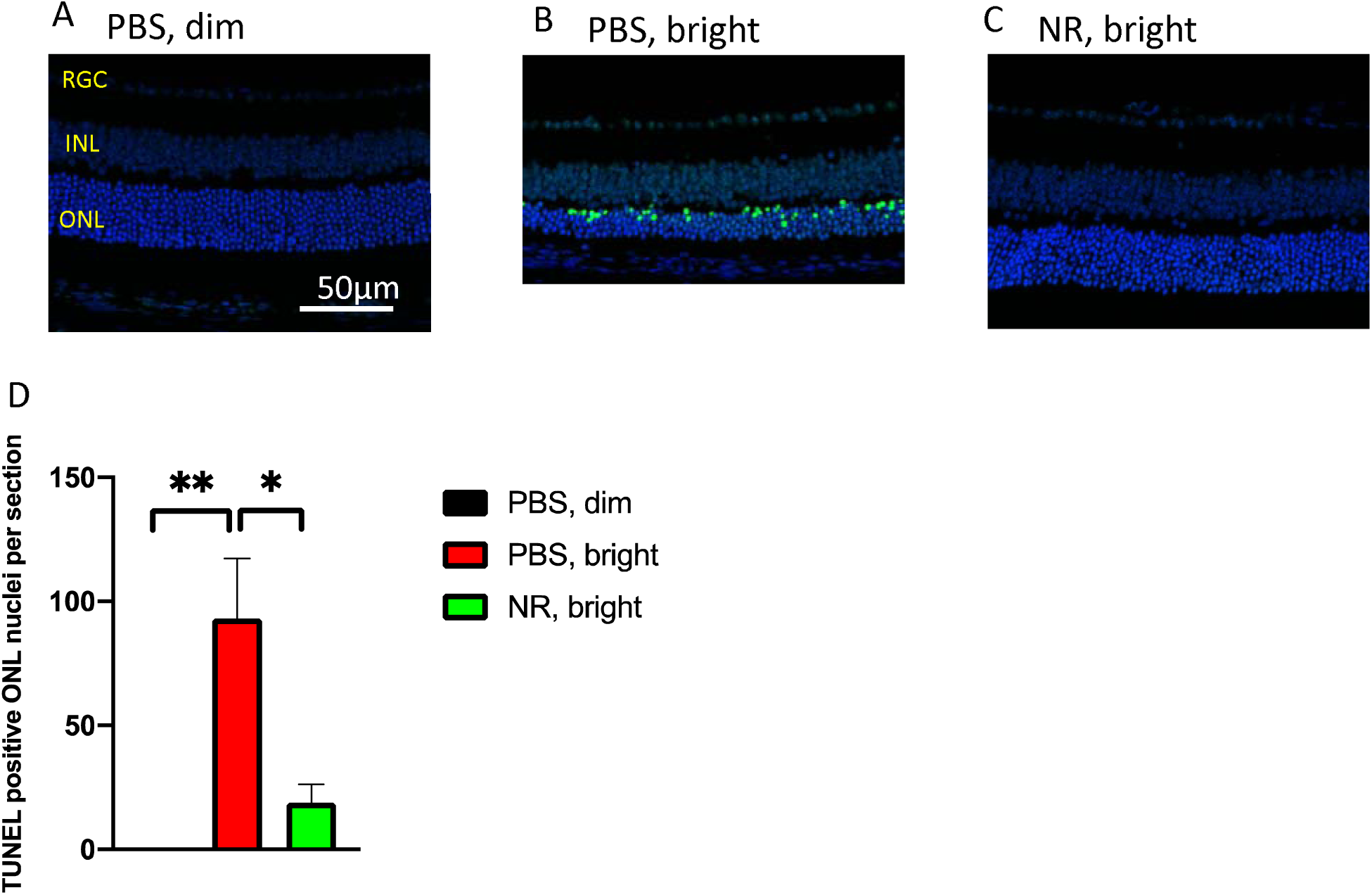
NR treatment prevents increases TUNEL-positive cells following induction of LIRD. Both PBS and NR treated Balb/c mice received toxic light exposure and were euthanized 1 week after exposure. Retinas from mice treated as described in text were fixed, sectioned, and used in a TUNEL assay. (A)-(C). Representative morphological images of each group. Complete sections are shown in Supplemental Figure S2. (D). TUNEL positive nuclei in ONL were counted from the entire retina. NR treated mice (red bar) exhibited significantly fewer TUNEL-positive cells compared to the PBS treated group (green bar). *P < 0.05, **P<0.01 by one-way ANOVA with Tukey’s multiple comparisons non-parametric test. n = 4 retinal images/group. Error bars represent SEM. Size marker represents 50µm.

### NR treatment prevents an inflammatory response following induction of LIRD

Subretinal autofluorescent spots observed *in vivo* by fundus examination in patients and in animal models are considered diagnostic markers for inflammatory responses in retinal damage and disease.^56–60^ To test whether NR treatment alters inflammatory responses of the LIRD mice, *in vivo* fundus examination at the level of the subretinal space was conducted one week following induction of retinal degeneration. Uninduced eyes showed few blue autofluorescent spots at the level of the subretinal space in vivo (Fig. 10A). In eyes of induced mice treated with PBS, numerous and widespread autofluorescent spots were observed (Fig. 10B). This was prevented in mice treated with NR, which exhibited fewer autofluorescent spots (Fig. 10C), similar in number and pattern to the uninduced group, confirmed by statistical testing on counts of these spots across several autofluorescent fundus images (Fig. 10D). These data suggest that LIRD leads to an inflammatory response that is largely prevented by NR treatment.

**Figure 10.**
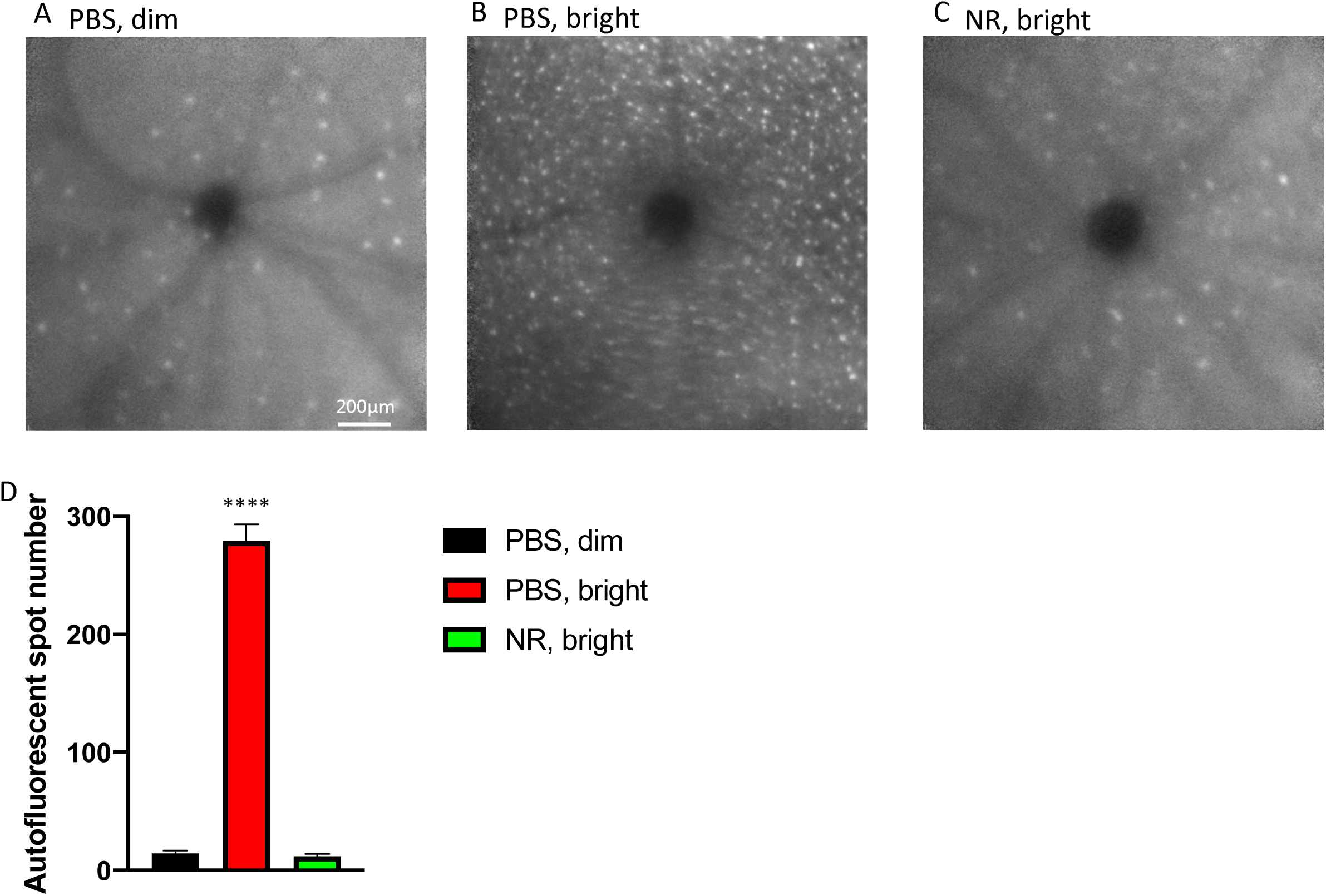
NR treatment prevented subretinal autofluorescence observed in vivo in LIRD mice. (A)-(C). Representative morphology image from each group at the level of the photoreceptor-RPE interface. In vivo Spectralis HRA+OCT images (with blue autofluorescence detection) were taken one week following induction of degeneration. (D). Autofluorescent white spots were counted across the fundus image FIELD. Few were detected in uninduced mice (black). Bright light exposed mice treated with NR exhibited significantly fewer white dots compared to the PBS treated group. ****P < 0.0001 one-way ANOVA with Tukey’s multiple comparisons non-parametric test. n = 3-6 mice/group. Error bars represent SEM. Size marker represents 200µm.

### NR elevates retinal NAD^+^ levels in mice

NR is a NAD^+^ precursor that when given systemically increases NAD^+^ levels in many tissues, including CNS structures,^29, 35, 36^ and its protective effects are ascribed in part to local NAD^+^ increases in target tissue.^30–32, 34, 36^ As an initial test of whether this may be the case in NR-induced retinal neuroprotection, we used C57Bl/6J mice to test whether NR treatment could increase NAD^+^ levels in retina. We IP-injected mice with either NR or PBS daily for 5 days. One hour after the last treatment, retinas were harvested and assayed for NAD^+^/NADH content, revealing that NR treatment increased retinal NAD^+^ (Fig 11A.) and NAD^+^/NADH ratio (Fig 11B) compared to the PBS-treated group.

**Figure 11.**
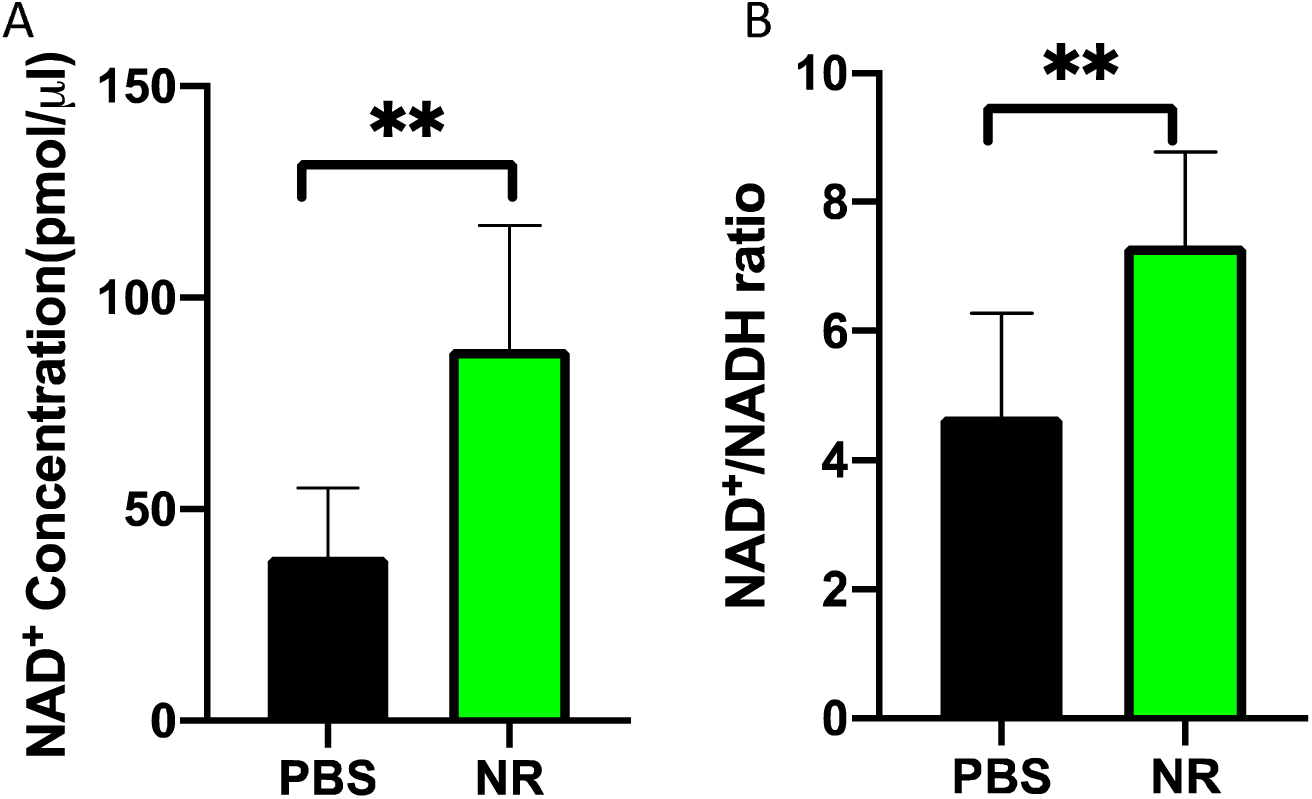
NR treatment increased retinal NAD^+^ levels in C57Bl/6J mice. Mice were IP injected with 1000 mg/kg NR or PBS for 5 consecutive days. Two hours after the last injection, they were sacrificed and retinas harvested. NAD^+^ and NAD(total) were assessed by colorimetric assay. Retinal NAD^+^ levels (A) and NAD^+^/NADH ratio (B) from mice treated with NR were statistically significantly increased compared to vehicle (PBS) treated group. **P<0.01 by Student *t*-test. n = 6 retinas per group. Error bars represent SEM.

## Discussion

In this study, we found that systemic treatment with NR protected photoreceptors from toxic light damage and delayed inherited retinal degeneration. NR has been shown to be protective in many models of neurodegenerative diseases, like Parkinson’s disease^35^ and Alzheimer’s disease.^29^ Our study is the first to demonstrate NR’s protective effects in retinal degeneration models.

Light-induced retinal damage is an acute model of retinal degeneration with a well-described oxidative stress response.^61^ We assessed by ERG the efficacy of NR treatment in preserving phoreceptor function in mice undergoing LIRD (Fig. 3). Remarkably, just two NR IP injections completely preserved scotopic a- and b-wave amplitudes. *In vivo* OCT imaging and fundus imaging showed that systemic NR treatment protected photoreceptors and whole retina from degeneration (Fig 7). Postmortem H&E staining provided more detailed information on retinal morphology, indicating that NR treatment protected retinal structure, both inferior and superior (Fig. 8). Retinas of mice undergoing LIRD exhibited marked TUNEL staining, especially in the ONL, suggesting massive apoptosis of photoreceptor cells (Fig. 9, compare Panel A with Panel B). NR treatment largely prevented this, suggesting that apoptosis was suppressed (Fig 9).

Autofluorescent spots similar to those observed here are postulated to be components of inflammatory responses, like activated microglial cells, lipofuscin in RPE cells, bisretinoids in the photoreceptors,^62–65^ and RPE cells undergoing epithelial-mesenchymal transition (EMT) that are migrating into the neural retina.^66, 67^ The present data indicate that NR treatment prevented the appearance of these autofluorescing entities (Fig 10), and thus, may be preventing retinal inflammation, possibly indirectly by partially preventing early rod photoreceptor injury, or more directly by some as-yet undefined action.

Although the LIRD mouse model of retinal degeneration has been extremely valuable in increasing our understanding of photoreceptor and retinal degeneration,^37, 68, 69^ it is a damage model and not a disease model. The rd10 mouse and the IRBP KO mouse are considered models of arRP.^42, 46, 47, 50, 70–76^ We found that retinal NR protection extends to these two arRP models (Fig 4-6), with significant increases of ERG amplitudes well after photoreceptor degeneration onset and retinal function loss in untreated cohorts.^42, 46, 47, 50, 70–74, 76^ The NR-induced increase in ERG mean amplitudes in IRBP KO mice, though significant (Fig. 4), was modest most likely because retinal function has not yet declined dramatically at this early stage in the degeneration.^40, 54^ The NR-induced increases in mean amplitudes of scotopic b-waves and photopic a- and b-waves in P28 rd10 mice were more dramatic and persisted at least 10 days after injections were halted (Figs. 5 and 6), demonstrating that NR treatment is capable of preserving cone photoreceptor and inner retinal function even at a stage at which extensive degeneration and function loss otherwise occurs.^42, 72, 75^ Rod photoreceptor function was not protected, unsurprising given that the mutation underlying the degeneration is in a rod-specific phototransduction gene.^42, 72^ Overall, it appears that systemic treatment with NR protects against both retinal damage and disease.

Photoreceptors and RPE have high metabolic demands.^15^ It may be that the observed increase in NAD^+^ and NAD^+^/NADH ratio following NR treatment (Fig. 11) allowed increased retinal metabolic capacity that contributed to protection.^77^ Recent work investigating the effects of NR in yeast and mammals established that its metabolic pathway involves kinases NMRK1 and NMRK2, which were first described by Brenner and colleagues in 2004.^27^ Apte and collegues demonstrated that an intact nicotinamide-NAMPT-NAD+ pathway is requisite for retinal health, but whether the same holds true for an NR-NMRK1/2-NAD+ pathway has not been explored. Additionally, it may be useful to compare the efficacy and potency between NR and nicotinamide treatment in retinal degeneration. Nicotinamide treatment has been well-investigated in neurodegenerative diseases, in mouse glaucoma models,^22, 23, 78, 79^ and in a rat LIRD model,^80^ but has yet to be test in a mouse retinal degeneration model.

In summary, this is the first study to demonstrate that systemic treatment with nicotinamide riboside is protective for photoreceptors undergoing stress in damage and in genetic degeneration. The protection is significant, and may support the proposition for prospective human subject studies.

## Supporting information

NR treatment preserves nuclei of the outer nuclear layer in the LIRD mice.

NR treatment prevents increases TUNEL-positive cells following induction of LIRD.

## ACKNOWLEDGMENTS

The studies were funded by the Abraham J. and Phyllis Katz Foundation (JHB); The joint training program between Emory University School of Medicine and Xiangya School of Medicine, Central South University. China Scholarship Council (201806370277 XZ); NIH R01EY028859 (JHB); NIH R01EY021592 (JMN); NIH R01EY028450 (JMN); VA I01RX002806 (JHB); VA I21RX001924 (JHB); VARR&D C9246C (Atlanta VAMC); NIH P30EY06360 (Emory); and an unrestricted departmental award from Research to Prevent Blindness. Inc. to the Ophthalmology Department at Emory University.

---

**Figure S1. NR treatment preserves nuclei of the outer nuclear layer in the LIRD mice**. A week after toxic light exposure, mice were euthanized and ocular sections stained with hematoxylin and eosin (H&E). Representative H&E images of complete retina sections from each treatment group are shown. **Panel A**: Section from a PBS-treated mouse exposed to dim light shows unperturbed ONL across the span of the retina. **Panel B**: Section from a PBS-treated mouse with toxic light exposure shows extreme thinning in the central, superior (right side of section) ONL. **Panel C**: Section from a NR-treated mouse exposed to toxic light shows similar thickness ONL as that in control mouse.

**Figure S2. NR treatment prevents increases TUNEL-positive cells following induction of LIRD**. A week after toxic light exposure the mice were euthanized and ocular sections stained for TUNEL (green) and DAPI (blue). Representative images of complete retina sections from each treatment group are shown. The right half of each section is dorsal/superior and the left half is ventral periphery/inferior. **Panel A**: Section from a PBS-treated mouse exposed to dim light shows little to no TUNEL-positive ONL nuclei across the span of the retina. **Panel B**: Section from a PBS-treated mouse with toxic light exposure shows numerous TUNEL-positive ONL nuclei in both superior and inferior halves. **Panel C**: Section from a NR-treated mouse exposed to toxic light shows much fewer TUNEL-positive ONL nuclei relative to PBS-treated mouse section.

