## Supplementary figures and images for "Systemic Treatment with Nicotinamide Riboside is Protective in Three Mouse Models of Retinal Degeneration"

### NR treatment prevents increases TUNEL-positive cells following induction of LIRD.

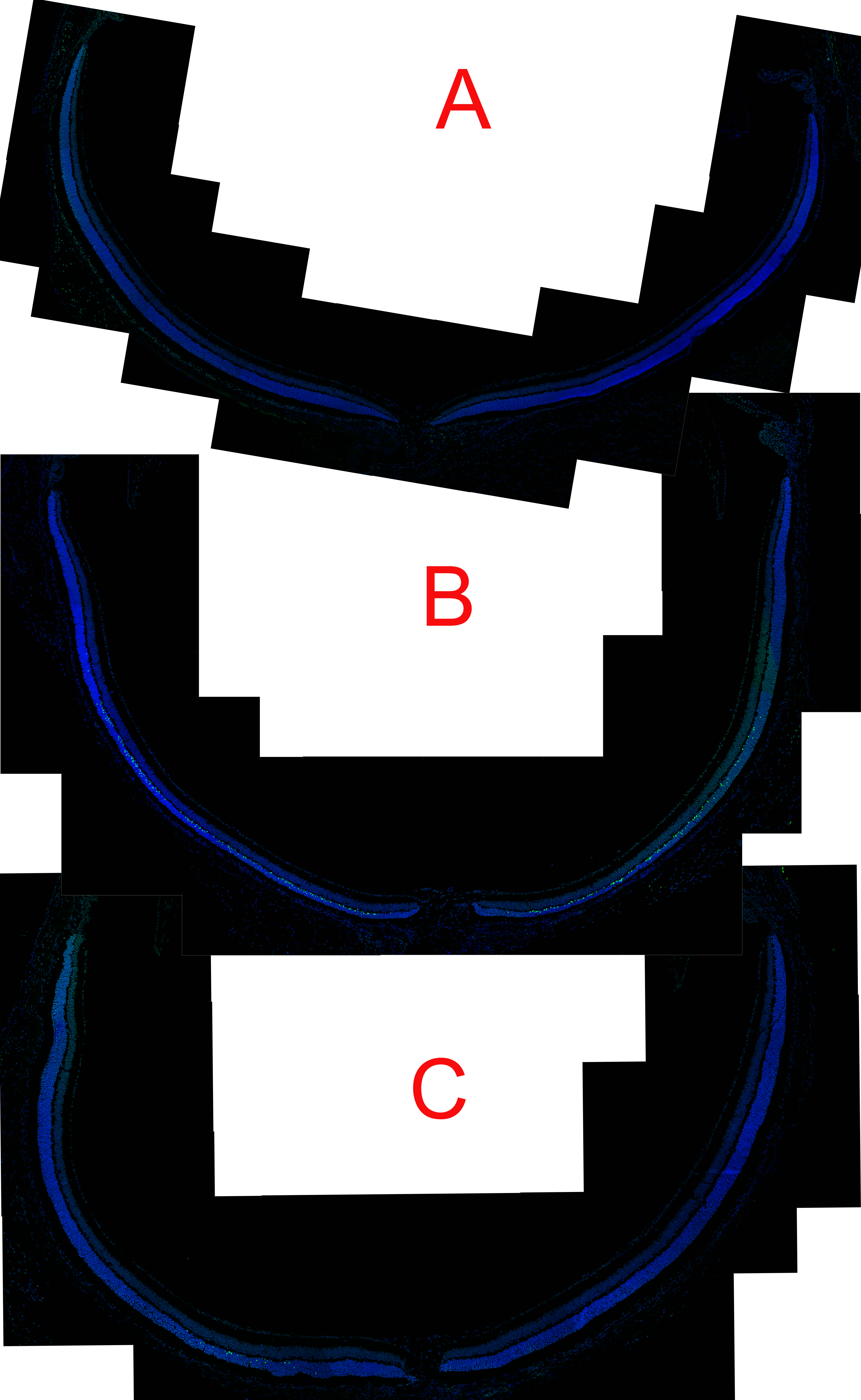
